# Gut microbiota-based vaccination engages innate immunity to improve blood glucose control in obese mice

**DOI:** 10.1101/2021.07.15.452537

**Authors:** Brittany M. Duggan, Akhilesh K. Tamrakar, Nicole G. Barra, Fernando F. Anhê, Gabriella Paniccia, Jessica G. Wallace, Hannah D. Stacey, Matthew S. Miller, Deborah M. Sloboda, Jonathan D. Schertzer

## Abstract

Obesity and diabetes increase circulating levels of microbial components derived from the gut microbiota. Individual bacterial factors (i.e., postbiotics) can have opposing effects on metabolic inflammation and blood glucose control. We tested the net effect of gut bacterial extracts on blood glucose using a microbiota-based vaccination strategy in mice. Male and female mice had improved insulin sensitivity and blood glucose control five weeks after a single subcutaneous injection of a specific dose of a bacterial extract obtained from the luminal contents of the proximal gut. Injection of mice with proximal gut extracts from germ-free mice revealed that bacteria were required for a microbiota-based vaccination to improve blood glucose control. Vaccination of *Nod1^−/−^*, *Nod2^−/−^*, and *Ripk2^−/−^* mice showed that each of these innate immune proteins was required for bacterial extract injection to improve blood glucose control. A microbiota-based vaccination promoted a proximal gut immunoglobulin-G (IgG) response directed against bacterial extract antigens, where subcutaneous injection of mice with the luminal contents of the ileum elicited a bacterial extract-specific IgG response that is compartmentalized to the ileum of vaccinated mice. A microbiota-based vaccination was associated with an altered the microbiota composition in the ileum and colon of mice. Lean mice required a single injection of proximal gut bacterial extracts, but high fat diet (HFD)-fed, obese mice required prime-boost bacterial extract injections for improvements in blood glucose control. These data show that, upon subversion of the gut barrier, vaccination with proximal gut bacterial extracts engages innate immunity to promote long-lasting improvements in blood glucose control in a dose-dependent manner.

## Introduction

The gut microbiota shapes host innate and adaptive immune responses, which reciprocally regulate the composition and function of the gut microbiota (Cox and Blaser, 2015; Gilbert et al., 2018; Surana and Kasper, 2014). This is relevant to metabolic disease because metabolic inflammation (i.e., metaflammation) contributes to insulin resistance and impaired blood glucose control (Hotamisligil, 2017). A common hypothesis is that obesity is associated with systemic inflammation. Obesity can coincide with low-level inflammation in metabolic tissues that regulate blood glucose, such as adipose and liver (Gregor and Hotamisligil, 2011; Kane and Lynch, 2019; Mathis, 2013). However, some compartmentalized immune responses in the proximal gut are lower in mouse models of obesity or prediabetes compared to their lean, normoglycemic counterparts (Garidou et al., 2015; McPhee and Schertzer, 2015). For example, there are fewer intestinal immunoglobulin A (IgA)-producing cells and less secretory IgA-associated immune mediators in high fat diet (HFD)-fed obese mice (Luck et al., 2019). There are also reductions in Cd4+ and Cd8+ T cells and impaired Th17 immune responses in the ileum of mice fed obesogenic and diabetogenic diets (Cavallari et al., 2016; Garidou et al., 2015; Hong et al., 2013). These data show that immunological changes during obesity are not restricted to chronically increased inflammation. In some circumstances, obesity is associated with a dampening of the immune responses that may be compartmentalized to specific tissues or cell populations.

The triggers of compartmentalized inflammation during obesity are largely unknown, although it has been proposed that changes in the gut microbiota (i.e., dysbiosis) promote metaflammation and consequently poor blood glucose control during obesity. Indeed, obesity is associated with changes in bacterial diversity and composition in rodents and humans and colonizing germ-free mice with obese microbiota promotes adiposity and poor blood glucose control (Ley et al., 2006, 2005; Turnbaugh et al., 2006). These data suggest a causal role for gut microbiota derived factors in metabolic and endocrine control (Foley et al., 2020, 2018; Ridaura et al., 2013; Turnbaugh et al., 2006). Consequently, altering the gut microbiota composition and fecal microbial transfer (FMT) have been proposed to mitigate insulin resistance and dysglycemia (Anhê, 2015; Kootte et al., 2017; Vrieze et al., 2012; Yoon et al., 2021). Which microbial strains are responsible for improved metabolic control is unclear, however specific commensal bacteria have been suggested to protect against aspects of metabolic disease. For example, intestinal abundance of *Bifidobacterium* species and *Akkermansia muciniphila* has been inversely associated with obesity, insulin resistance and type 2 diabetes (T2D) (Depommier et al., 2019; Everard et al., 2013; Liou et al., 2013; Million et al., 2012; Pedret et al., 2019; Plovier et al., 2016). It is still not clear how the multiple functions of gut microbe communities or specific bacterial strains can be harnessed to alter blood glucose, but emerging evidence shows that gut microbes impact host health through changes in circulating metabolites (Chen et al., 2021).

Bacteria and their components or metabolites can subvert the intestinal barrier and reach extra-intestinal tissues to exert compartmentalized immune and metabolic responses (Anhê et al., 2020; Massier et al., 2020). Increased systemic bacterial load is proposed to worsen blood glucose control, where T2D alters bacterial DNA signatures and indicators of live bacteria in adipose tissue of humans with obesity (Anhê et al., 2020; Massier et al., 2020). However, live bacteria are not required to alter host immunity and metabolism. Bacterial-derived factors, cell components and metabolites, collectively termed postbiotics, can alter host blood glucose homeostasis (Anhê et al., 2021). For example, subverting the gut barrier by injection or infusion with specific bacterial cell-wall components such lipopolysaccharide (LPS) and meso-diaminopimelic acid (meso-DAP)-containing muropeptides triggers inflammatory responses and exacerbates insulin resistance and dysglycemia in mice (Cani et al., 2007; Duggan et al., 2017; Schertzer and Klip, 2011). Conversely, repeated injection of the bacterial cell wall component muramyl dipeptide (MDP) lowers adipose tissue inflammation and mitigates insulin resistance in obese mice by engaging a Nucleotide Binding Oligomerization Domain Containing 2 (NOD2) innate immune response (Cavallari et al., 2020a, 2017). Single bacteria factors can have opposing effects on blood glucose, but the net effect of various systemic postbiotics derived from the gut microbiota is not yet clear.

Bacteria can alter immune responses beyond acute pro-inflammatory or anti-inflammatory actions. Immunological memory allows for a rapid and effective response upon secondary encounter with the same antigen through adaptive immunity (Ratajczak et al., 2018). Innate immune cell memory responses are emerging, but poorly defined in metabolic disease (Bekkering et al., 2020; Penkov et al., 2019; Yao et al., 2018). A seminal study showed that subcutaneous injection of a crude total extract derived from luminal contents in the ileum of mice, generated an adaptive immune response that partially protected mice from developing glucose intolerance and insulin resistance (Pomié et al., 2016). This “vaccination-like” strategy provided long-lasting improvements in blood glucose control when used prior to feeding mice a diabetogenic diet. Improved glycemia required an adaptive immune system, since *Rag1^−/−^* mice lacking lymphocytes were refractory to vaccination-induced effects on glucose tolerance, and adoptive transfer of immune cells from bacterial-vaccinated mice into naïve mice protected against dysglycemia induced by a diabetogenic diet in mice (Pomié et al., 2016). The previous study used a diabetogenic diet that did not induce obesity and the role of the innate immune system was not investigated. Here, we tested the net effect of postbiotics found in the gut microbiota on immune responses and blood glucose regulation in lean and diet-induced obese mice, using a microbiota-based vaccination strategy. We found that subcutaneous injection of proximal gut bacterial luminal extract elicited an IgG response in the ileum, altered the composition on the gut microbiome and promoted long-lasting improvements in blood glucose control. We found that a microbiota-based vaccination strategy required a NOD2-mediated innate immune response and that obese mice required prime-boost injections to improve glucose control.

## Methods

### Mice & materials

All animal procedures for this study were approved by the McMaster University Animal Research Ethics Board in accordance with the guidelines of the Canadian Council of Animal Care. All mice were a minimum of 8 weeks old before experiment initiation. Mice were maintained on a 12-hour light/dark cycle, and experiments were performed on multiple cohorts of mice born from different parents at different times of the year. WT (C57BL/6J), *Nod1−/−*, *Nod2−/−* and *Ripk2−/−* mice used for experiments were bred in-house under specific pathogen-free conditions at McMaster University. Germ-free mice were obtained from the Farncombe Gnotobiotic Unit of McMaster University. All animals were fed a control diet (CD) (17% kcal from fat, 29% kcal from protein, 54% kcal from carbohydrate: cat# 8640 Teklad 22/5, Envigo). Where indicated, mice were fed an obesity-promoting 60% HFD (60% kcal from fat, 20% kcal from protein, 20% kcal from carbohydrate: cat# D12492). Glucose tolerance tests (GTTs) and insulin tolerance tests (ITTs) were performed in 6 h-fasted, conscious mice and oral glucose-stimulated insulin secretion tests (OGSIS) were performed in 12 h-fasted, conscious mice (Denou et al., 2015). The dose of D-glucose (Sigma-Aldrich) or insulin (NovoRapid, Novo Nordisk) and route of administration using intraperitoneal (i.p.) injection or oral gavage (p.o.) is indicated in each figure caption for each experiment. Blood glucose was determined by tail vein blood sampling using a handheld glucometer (Roche Accu-Check Performa). During OGSIS tests, blood samples were collected via tail-vein sampling at time (t)=0-, 10- and 60-min post-glucose gavage (4g/kg). Blood was centrifuged for 10 min at 4°C and 10,000 g, and plasma fraction was collected and stored at −80°C. Plasma insulin levels were assessed using high sensitivity mouse insulin ELISA kit (Toronto Bioscience, Cat# 32270) and measured with a Synergy H4 Hybrid reader (Biotek Instruments). Area under the curve (AUC) of blood glucose or blood insulin vs. time was calculated for GTT, ITT and OGSIS using GraphPad Prism 6 software (with baseline Y values set to 0).

### Collection and preparation of intestinal extracts for vaccination

Each intestinal extract contained the luminal contents of 3 donor mice of the same sex as the recipient mice. Groups of 3 WT/J donor mice, aged 10-14 weeks, were used for preparation of intestinal extracts and mice were fed a control (CD) or high fat diet (HFD) for 4 weeks prior to euthanization via cervical dislocation. Subsequently, the entire length of the both the small and large intestine were removed with sterile tools. Luminal contents of each intestinal section were collected via gentle and thorough pressurization into a clean tube containing 500μL DPBS on ice, using separate tools for each section. The duodenal/jejunal section was defined as the proximal 15cm of small intestine, measured distally from the pyloric sphincter. The ileal section was defined as the distal 10 cm of small intestine, starting from cecum and measuring proximally. The entire contents of the cecal sack were collected for cecum extracts. The entire length of colon from cecum to rectum was collected for colon extracts. Upon collection, all luminal contents were vigorously vortexed for 1 min to ensure thorough mixing. For fecal extracts, a single fecal pellet was collected from each of the 3 donor mice prior to euthanization, and mechanically homogenized at 4.5 m/s for 1 min using a FastPrep-24 tissue homogenizer (MP Biomedicals) and two plastic beads. Germ-free intestinal extracts were prepared from 3 age-matched germ-free mice immediately upon export from the Farncombe Gnotobiotic Unit at McMaster University. Euthanization and intestinal extracts were prepared in a level II biosafety hood to limit contamination with ambient microbes. All extracts were centrifuged (4°C, 7500rpm, 5 min) to pellet debris and supernatant was transferred into fresh tubes. Supernatant was sonicated for 1 min (Fisher Scientific 20kHx sonicator), aliquoted and stored at −80° until use). Extracts were stored for a maximum of 4 months, and multiple separate collections of extracts were tested across multiple cohorts of mice. The pooled intestinal extract from 3 mice was combined and used to inject (i.e., vaccinate) multiple mice of the same sex for each experiment.

### Microbiota-based vaccination

Luminal contents from the ileum were used unless otherwise indicated. Diluted extracts from the ileum (or each gut segment) were prepared fresh on the day of injection in recipient mice in dPBS (50-20,000x dilution). WT (C57BL/6J, male and female) and *Nod1−/−*, *Nod2−/−* and *Ripk2−/−* male mice, aged 10-18 weeks old, received a single subcutaneous injection of intestinal extract (200μL, *s.c.*, of duodenal/jejunal, ileal, cecal, colon or fecal extract, as indicated). Glucose tolerance, insulin tolerance or oral glucose-stimulated insulin secretion was assessed 4-5 weeks after injection, as indicated.

For the “prime-boost” vaccination model, male mice received a first injection of ileal extract (200μL, *s.c.*) after 8 weeks of HFD-feeding, and a second injection was administered after 12 weeks of HFD-feeding. Glucose tolerance was assessed after one injection at 12 weeks and two injections at 16 weeks of HFD-feeding. In the CD-to-HFD prime-boost model, male mice were fed a CD, injected once and glucose tolerance was assessed after one injection at 5 weeks of CD feeding. Subsequently, 5 weeks after the initial injection, CD mice were switched to a 60% HFD. After 8 weeks on a HFD mice were injected a second time. Glucose tolerance was tested 4 weeks after the secondary ‘boost’ injection after a total of 12 weeks of HFD-feeding.

### Quantification of IgG in serum & intestinal samples

Serum, ileum, cecal, and colon samples were harvested from vaccinated male mice 6 weeks after the vaccination event. Whole blood was collected via facial vein sampling in conscious mice. Mice were then immediately euthanized by cervical dislocation and intestinal segments were immediately excised. Blood was clotted for 20 min and then centrifuged for 10 min at 4°C and 10,000 x g. The serum fraction was collected and stored at −80° until assay. Intestinal samples (ileum, cecum, colon) were homogenized in 1.5 ml Rino Tubes (Next Advance, Troy, NY, USA) containing 1.6 mm stainless steel beads (Next Advance, Troy, NY, USA) and 500µL dPBS using the Bullet Blender Gold (Next Advance) at maximum speed for 10 minutes at 4 °C and then centrifuged for 5 minutes at 8,000 g. Supernatants were collected and stored at −80° until assayed. Protein concentration of ileal extract was determined by Pierce BCA Protein assay (Thermo Scientific, Waltham, MA, USA,) according to manufacturer’s instructions and 96-well NUNC- Maxisorp plates (Thermo Scientific, Waltham, MA, USA,) were coated overnight at 4 °C with 2 μg/mL of the same ileal extract used for vaccination. The extract was diluted in bicarbonate-carbonate coating buffer (pH 9.4). The next day antigen-coating buffer was discarded, and plates were blocked by shaking for 1 hour at 37°C with reagent diluent (0.5% bovine serum albumin (BSA), 0.02% sodium azide, in 1X Tris-Tween wash buffer −10X composed of 0.024% Tris, 0.0876% Sodium Chloride, 0.00373% Potassium Chloride, 0.005% Tween-20 in 800ml MilliQ, adjusted to pH 7.4 with 3M HCl, final volume 1000ml). After blocking, samples were serially diluted from a 1:10 (serum) or 1:2 starting dilution (ileum, cecum, colon). Samples were incubated for one hour, shaking at 37°C. Following sample incubation, plates were washed three times with 1X Tris-Tween wash buffer. A goat anti-mouse IgG- biotin antibody (Southern Biotech) was diluted 1:5000 in reagent diluent and added to all wells. Plates were again incubated for 1 hour at 37°C with shaking, followed by three washes with 1X Tris-Tween buffer. A streptavidin-alkaline phosphatase secondary antibody (Southern Biotech) was diluted 1:2000 in reagent diluent and added to all wells followed by another 1-hour incubation at 37°C with shaking. Following this incubation, 3 more washes with 1X Tris-Tween buffer were performed, the final wash was removed and pNPP one component microwell substrate solution (Southern Biotech) was added to each well. After 10 minutes of developing, the reaction was quenched with 3N sodium hydroxide. The optical density (O.D.) at 410 nm was read on a Spectramax I3 (Molecular Devices, San Jose, CA, USA). IgG endpoint titers were defined by the lowest dilution at which the O.D. was three standard deviations above the mean of the blank wells.

### Bacterial profiling

Isolation of DNA from ileal and colon contents was done using mechanical and enzymatic lysis (Denou et al., 2016). The V3-4 region of the 16S rRNA gene was PCR amplified with barcode tags compatible with Illumina technologies and the Illumina MiSeq platform was used to sequence amplified DNA products. Details are available at www.surettelab.ca/protocols. A custom pipeline was used to process the FASTQ files. Dada25 (Callahan et al., 2016) was used to assign reads to Amplicon Sequence Variants (ASVs) and assign taxonomy with the Ribosomal Database Project (RDP) Bayesian classifier using the Silva 132 database (Quast et al., 2012). ASV assignments were converted to relative abundance before β-diversity calculations to account for depth of coverage and to normalize across samples. QIIME and R scripts were used to calculate β-diversity and to perform statistical tests.

### Statistical analysis

Individual data points indicate separate mice and data is also expressed as mean ± standard error of the mean (SEM). Comparisons were made using unpaired, two-tailed Student’s t-test where two variables are compared. ANOVA was used for comparison of more than 2 variables and Tukey’s post-hoc test was used when appropriate. Analysis of microbial populations was conducted in R. Partitioning of the variance in the microbiota was done with a Permutational multivariate analysis of variance (PERMANOVA) on Bray-Curtis dissimilarities calculated from relative amplicon sequence variant (ASV) abundance. The Wilcoxon rank sum test was used for pairwise comparisons. Adjustment for the false discovery rate (FDR) was calculated with the Benjamini-Hochberg method. R packages used for data analysis and visualization included vegan, ggplot2, tidyr, dplyr, ggtree, and corrplot. Significance was accepted at *p* < 0.05. Graphpad Prism 6-9 software was used for all analysis and diagrams were created using BioRender.com software. All data and R scripts generated in this study are available upon reasonable request.

## Results

### Microbiota-based vaccination with proximal-gut bacterial components improves blood glucose control in lean male mice

The luminal contents from the ileum were collected, sonicated and pooled from a group of 3 wild type C57Bl/6J male mice fed a control diet (CD). These ileum extracts were used for future subcutaneous injection (i.e., vaccination) into recipient male mice. Male mice fed a CD were vaccinated with a single injection of ileal extract 5 weeks before metabolic testing (Figure 1A). We found that injection of ileal extracts caused a dose-dependent change in blood glucose during a glucose tolerance test (GTT) but did not alter body mass (Figure 1B). Diluting ileal extracts by 5000x lowered blood glucose levels during a GTT, compared to less dilute (50x, 500x) and more dilute (20,000x) preparations of extract, demonstrating that a critical concentration of ileal extract was required to alter glycemia. This concentration of intestinal extract required for lowering blood glucose is consistent with published results (Pomié et al., 2016) and was used for all subsequent tests. Next, we found that injection of ileal extracts (5000x dilution) lowered blood glucose levels during an insulin tolerance test (ITT) (Figure 1C) but did not alter insulin secretion during an oral glucose load (Figure 1D).

**Figure 1.**
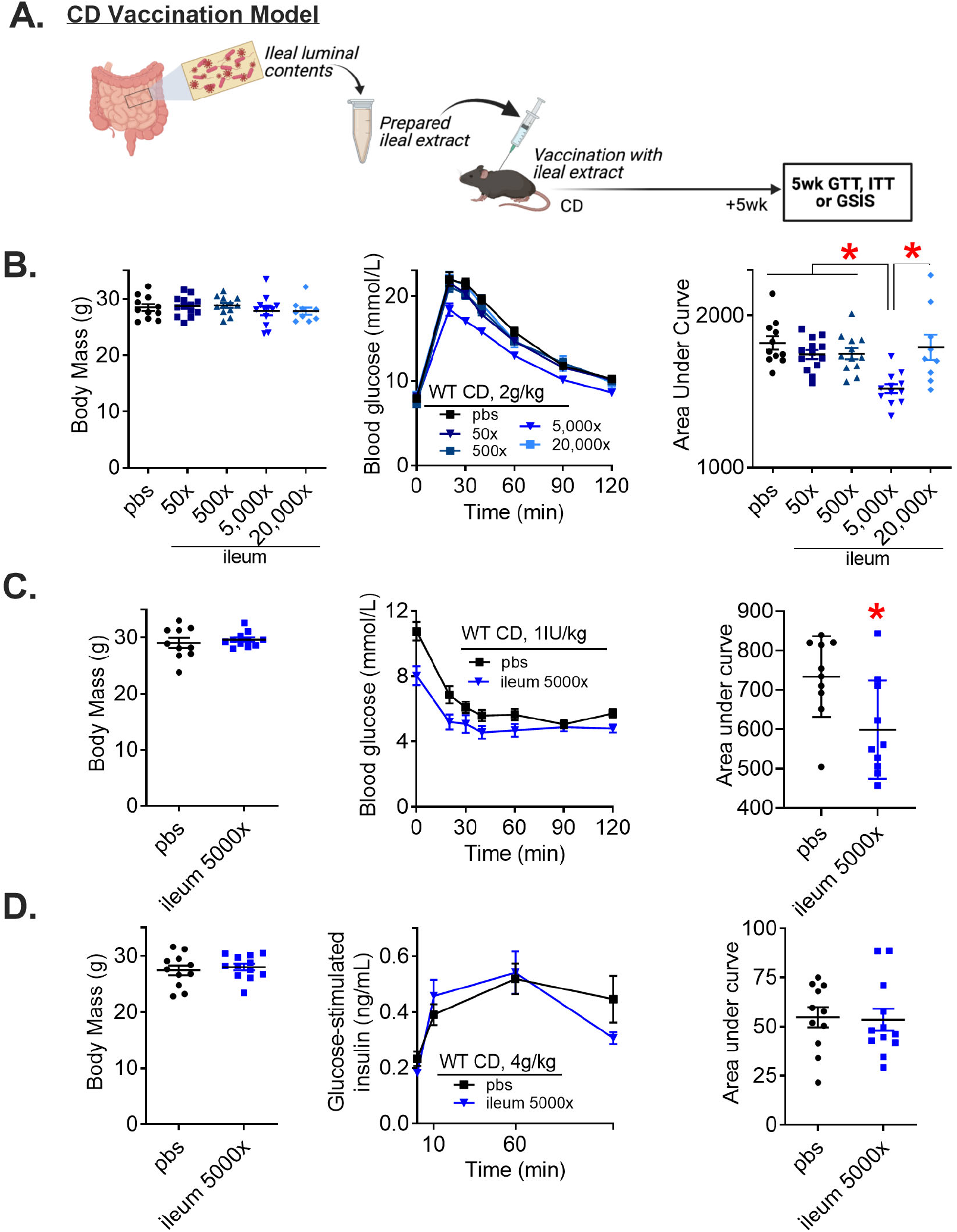
Subcutaneous injection of a specific concentration of ileal intestinal contents improves glucose and insulin tolerance in lean male mice. A) Experimental vaccination model of control diet (CD)-fed mice given a single subcutaneous injection with a previously prepared ileal extract. Glucose tolerance, insulin tolerance, and glucose-stimulated insulin secretion were assessed 5 weeks after vaccination. B) Body mass, blood glucose vs. time and quantified area under the curve during a glucose tolerance test (GTT) (2g/kg, i.p.) in CD-fed WT male mice 5 weeks after vaccination with various dilutions (50x-20,000x) of ileal extract (n=9-13). C) Body mass, blood glucose vs. time and quantified area under the curve during an ITT (1 lU/kg, i.p.) in CD-fed WT male mice 5 weeks after vaccination with a 5000x dilution of ileal extract (n=10-11). D) Body mass, plasma insulin concentration vs. time and quantified area under the curve during a GSIS (4g/kg, p.o.) (n=11-12). ‘Denotes significant different from control group injected with saline (pbs) determined by t-test or one-way ANOVA, where appropriate (p < 0.05). Each symbol represents a mouse and other values shown are the mean +/− SEM.

We postulated that the antigen(s) responsible for the glucose-lowering effects of ileal extracts were derived from bacteria. To determine if microbial components present in the ileal extracts were required for blood glucose-lowering, we next prepared ileal extracts from germ-free (GF) mice, and then vaccinated a cohort of CD-fed male mice (Figure 2A). Vaccination with ileal extract from male GF mice did not alter body mass or blood glucose levels in CD-fed male mice (Figure 2B). Thus, the glucose-lowering effects of ileal extracts are mediated by microbial components present in the intestinal milieu. This is consistent with published results that also demonstrated ileal extracts prepared from antibiotic-treated mice and then used for vaccination, did not alter glycemic control (Pomié et al., 2016). Finally, we tested the microbiota vaccination procedure using extracts harvested from different intestinal segments spanning the length of the small and large intestine in mice. In addition to the ileum, only extracts collected from the luminal contents of the duodenal/jejunal sections of the small intestine lowered blood glucose levels during a GTT, and this occurred independent of changes in body mass (Supplemental Figure 1). Taken together, these data show that proximal gut segments (i.e., duodenum, jejunum, and ileum) contain microbial factors (i.e., postbiotics) that improve glycemic control in lean male mice after a single vaccine-like subcutaneous injection.

**Figure 2.**
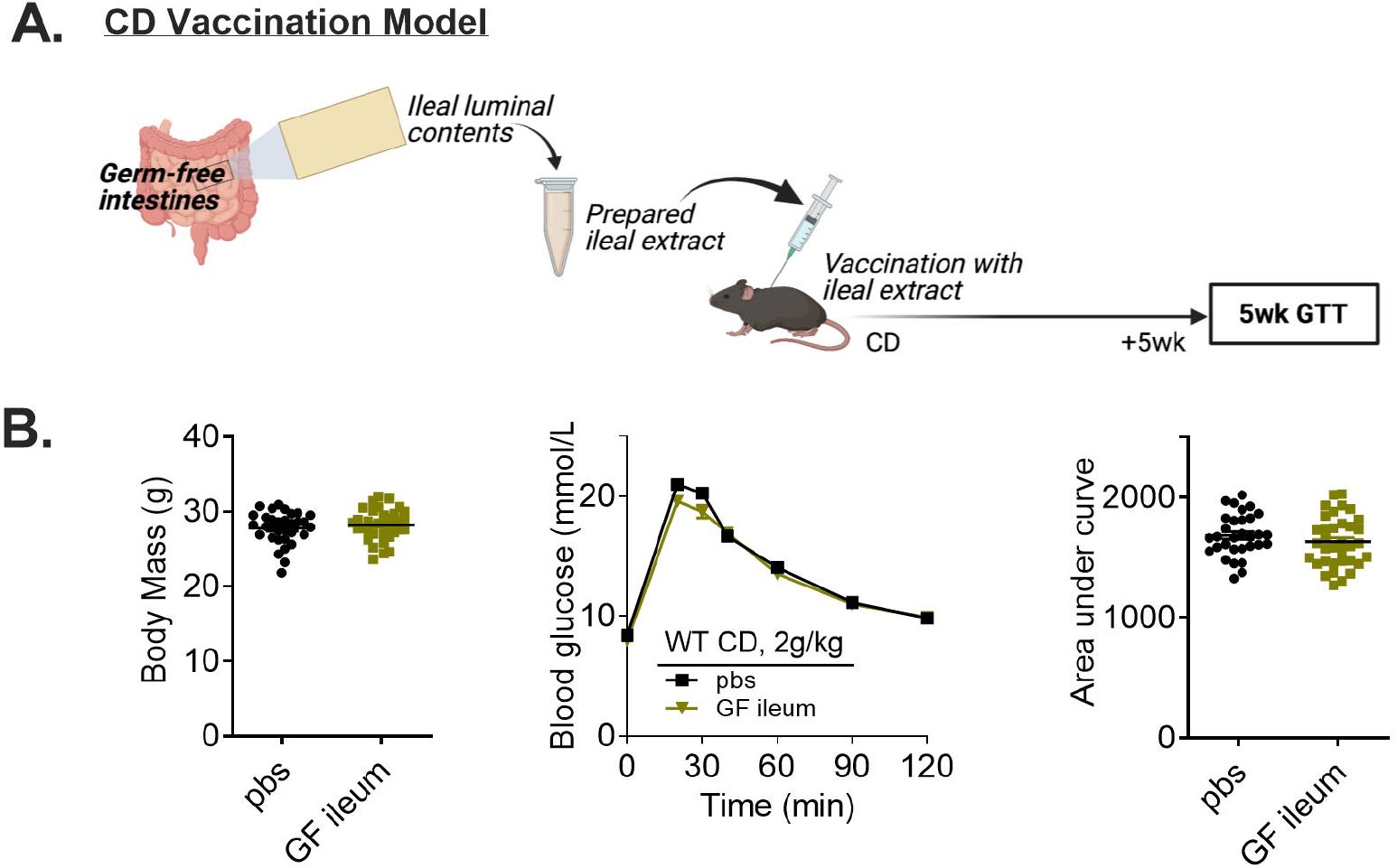
Bacteria are required fora microbiota-based vaccination to improve blood glucose control in lean male mice. A) Experimental vaccination model of control diet (CD)-fed mice given a single subcutaneous injection with an extract prepared from the ileal luminal contents of germ-free mice. Glucose tolerance was assessed 5 weeks after the vaccination event. B) Body mass, blood glucose vs. time and quantified area under the curve during a glucose tolerance test (GTT) (2g/kg, i.p.) in CD-fed WT male mice 5 weeks after vaccination with germ-free ileal extract (n=31-35). ‘Denotes significant different from control (pbs) group determined by unpaired t-test (p < 0.05). Each symbol represents a mouse and other values shown are the mean +/− SEM.

### Microbiota-based vaccination with bacterial extracts derived from both lean and obese mice improve glycemic control and effects are independent of sex

Diet is a major factor altering composition of the microbiota and even short-term changes in the relative amounts of ingested macronutrients produces rapid and distinct shifts in the microbial populations (Turnbaugh et al., 2009). Since sex also alters the composition of the gut microbiota (Org et al., 2016), we next tested whether an interaction exists between diet and sex to influence glycemic control after a microbiota-based vaccination. We used ileal extracts derived from male and female mice fed a CD or HFD and measured glycemia in mice vaccinated with ileal contents from the same sex. (Figure 3A). We found that ileal extracts from male and female mice fed a CD or HFD lowered blood glucose during a GTT in lean male and female mice without changes in body mass (Figure 3B, C). Taken together, our results show that postbiotics derived from the proximal gut of lean and obese mice (fed a HFD) possess blood glucose-lowering properties in both male and female mice.

**Figure 3.**
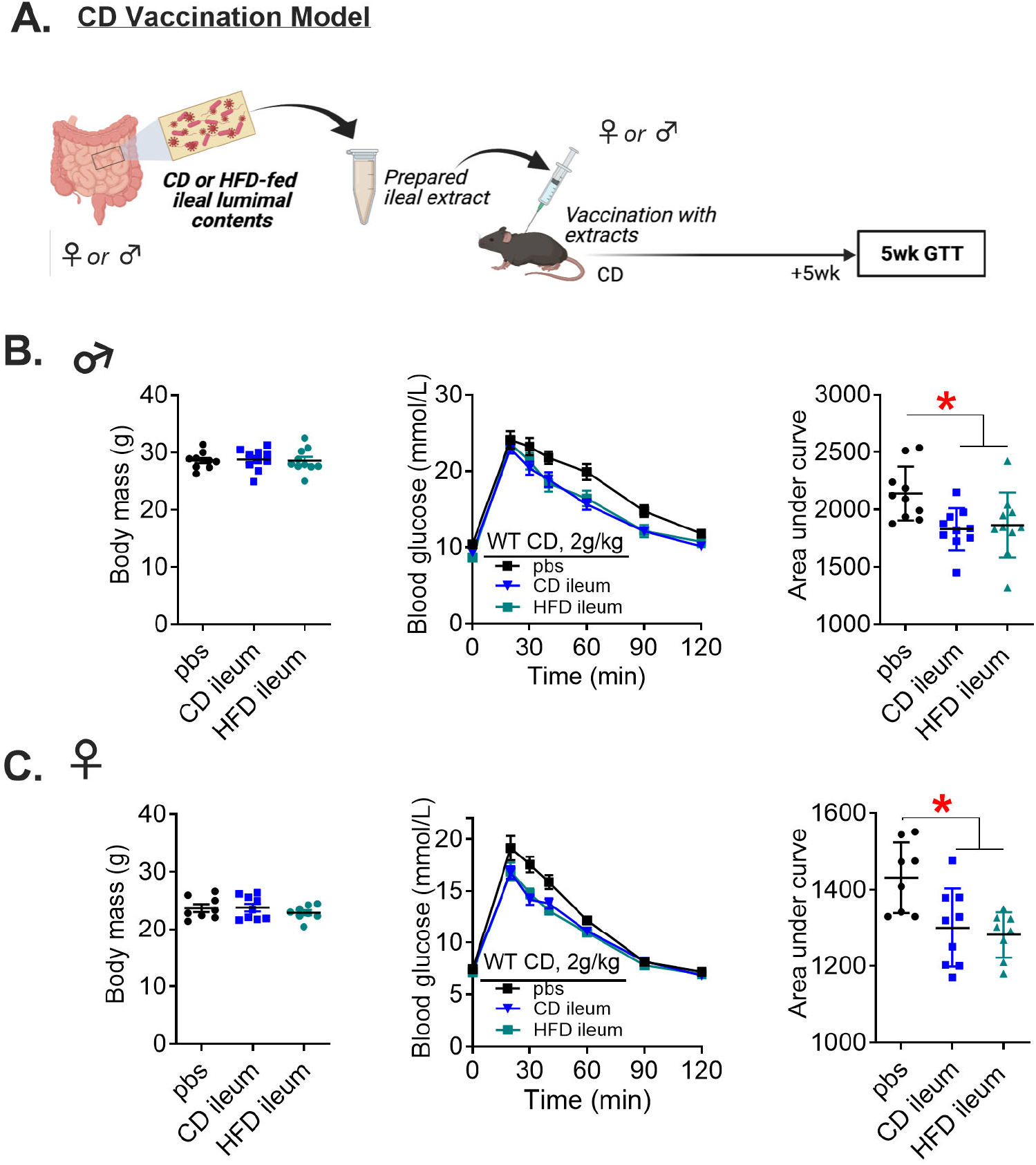
Microbiota-based vaccination with bacterial extracts derived from lean or obese mice improves blood glucose control in lean mice independent of sex. A) Experimental vaccination model of control diet (CD)-fed male mice given a single subcutaneous injection with an extract prepared from the ileal luminal contents of CD- or high fat diet (HFD)-fed male mice. Female CD-fed mice were given a single subcutaneous injection with an extract prepared from the ileal luminal contents of CD- or HFD-fed female mice. Glucose tolerance was assessed in all vehicle-injected and ileal-vaccinated mice 5 weeks after the vaccination event. B) Body mass, blood glucose vs. time and quantified area under the curve during a glucose tolerance test (GTT) (2g/kg, i.p.) in CD-fed WT male mice 5 weeks after vaccination with CD ileal extract or HFD ileal extract (n=10). C) Body mass, blood glucose vs. time and quantified area under the curve during a GTT (2g/kg, i.p.) in CD-fed WT female mice 5 weeks after vaccination with CD ileal extract or HFD ileal extract (n=8-9). ‘Denotes significant different from control group injected with saline (pbs) determined by one-way ANOVA (p < 0.05). Each symbol represents a mouse and other values shown are the mean +/− SEM.

### Microbiota-based vaccination engages NOD2 immunity to alter glycemic control

T-cell mediated adaptive immunity is required for improved glucose control after a bacterial vaccination strategy in mice (Pomié et al., 2016). However, it was not known if innate immune components are required. We tested if the bacterial cell wall sensors NOD1 and NOD2 and a common adaptor Receptor-interacting serine/threonine-protein kinase 2 (RIPK2) propagate changes in glycemia after bacterial vaccination in mice. CD fed wild type (WT), *Nod1^−/−^, Nod2^−/−^,* and *Ripk2^−/−^ male* mice were vaccinated with ileal extracts from CD-fed WT male mice. Our results confirm that WT mice have significantly lower blood glucose levels during a GTT five weeks after vaccination (Figure 4A). Unexpectedly, we found that *Nod1^−/−^* mice had higher blood glucose during a GTT five weeks after vaccination (Figure 4B). We found that *Nod2^−/−^* mice had no change in blood glucose during a GTT five weeks after vaccination (Figure 4C). *Ripk2^−/−^* mice had higher blood glucose during a GTT five weeks after vaccination (Figure 4D), which paralleled the blood glucose response in *Nod1^−/−^* mice after vaccination. Taken together, these results show that a peptidoglycan sensing NOD-RIPK2 mediated innate immune pathway is required for a microbiota vaccination strategy to improve blood glucose control in male mice. We found NOD2 is required for bacterial vaccination strategy to alter blood glucose in mice, whereas deletion of *Nod1* or *Ripk2* worsens glycemic control when male mice are injected with bacterial extracts.

**Figure 4.**
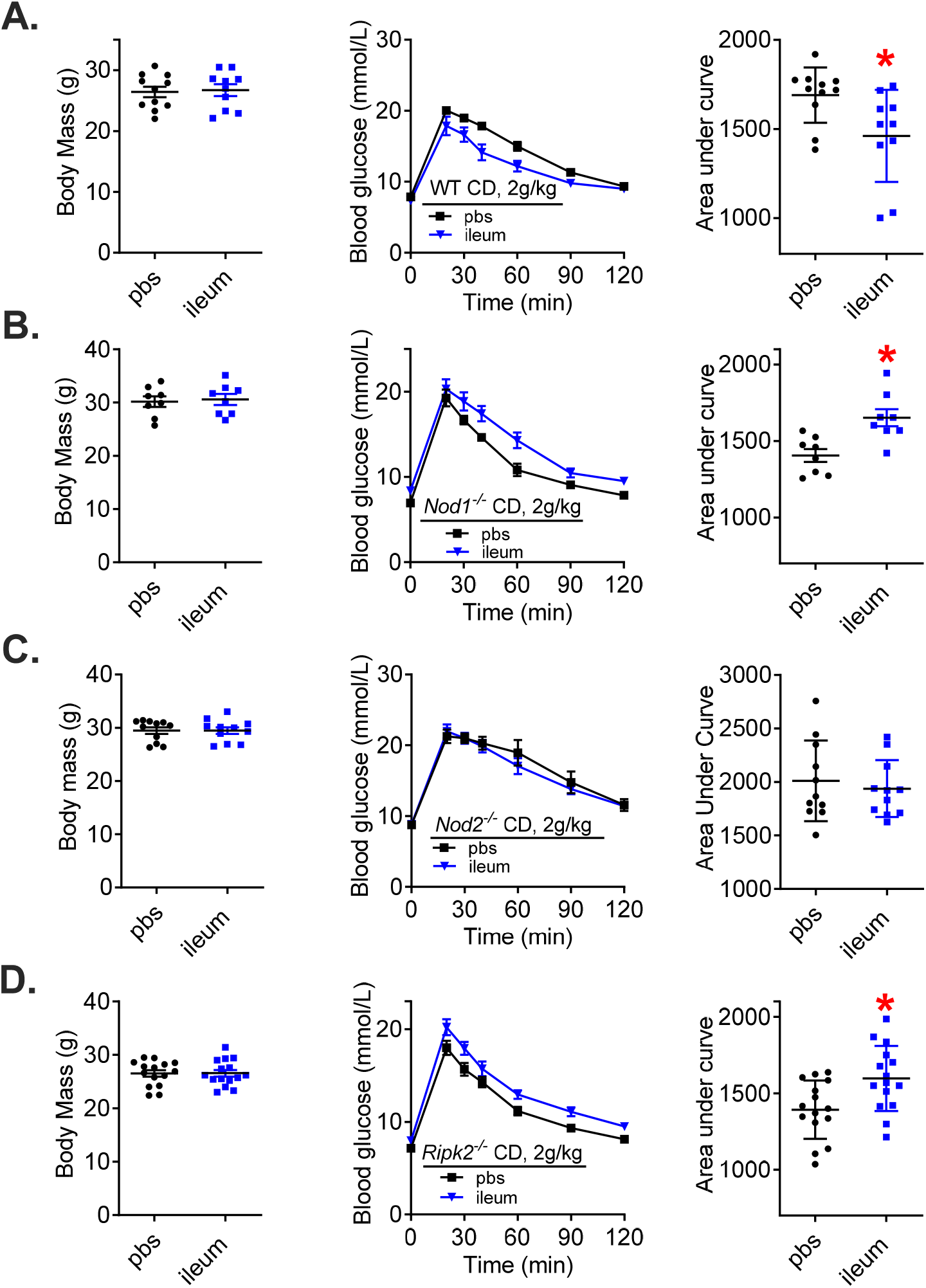
Microbiota-based vaccination engages NOD-RIPK2 to alter glycemia in lean male mice. A) Body mass, blood glucose vs. time and quantified area under the curve during a glucose tolearnace test (GTT) (2g/kg, i.p.) in control diet (CD)-fed wild type (WT) male mice 5 weeks after vaccination (n=10-11). B) Body mass, blood glucose vs. time and quantified area under the curve during a GTT (2g/kg, i.p.) in CD-fed Nod1^-/-^ male mice 5 weeks after vaccination (n=8). C) Body mass, blood glucose vs. time and quantified area under the curve during a GTT (2g/kg, i.p.) in CD-fed Nod2^-/-^ male mice 5 weeks after vaccination (n=11). D) Body mass, blood glucose vs. time and quantified area under the curve during a GTT (2g/kg, i.p.) in CD-fed Ripk2^-/-^ male mice 5 weeks after vaccination (n=15). *Denotes significant different from control group injected with saline (pbs) determined by t-test (p < 0.05). Each symbol represents a mouse and other values shown are the mean +/− SEM.

### A prime-boost microbiota-based vaccination improves glucose control in obese male mice

We next sought to determine if diet-induced obesity altered the efficacy of microbiota-based vaccination on glycemic control. Male mice were fed a HFD for 8 weeks and then injected with bacterial ileal extracts (Figure 5A). This initial “prime” vaccination did not alter body mass or blood glucose control 4 weeks later when the mice had been on a HFD for a total of 12 weeks (Figure 5B). These mice were then given a second injection with ileal extracts (Figure 5A). Four weeks after the subsequent “boost” vaccination, mice had lower blood glucose levels during a GTT, despite no change in body mass (Figure 5C). We then tested if the initial prime vaccination could be given in lean CD fed mice, followed the second boost vaccination in the same mice after HFD-feeding (Figure 5D). This prime-boost vaccination spanning a change from CD to HFD feeding lowered blood glucose levels during a GTT without a change in body mass (Figure 5E). Taken together, these data show that a ‘prime-boost’ vaccination strategy with ileal bacterial extracts is needed to improve blood glucose control during diet-induced obesity in male mice. A single injection of bacterial extracts was insufficient to improve blood glucose control in obese mice. Blood glucose levels could be lowered if the first (prime) and second (boost) injection were given to mice before and after the onset of diet-induced obesity, respectively.

**Figure 5.**
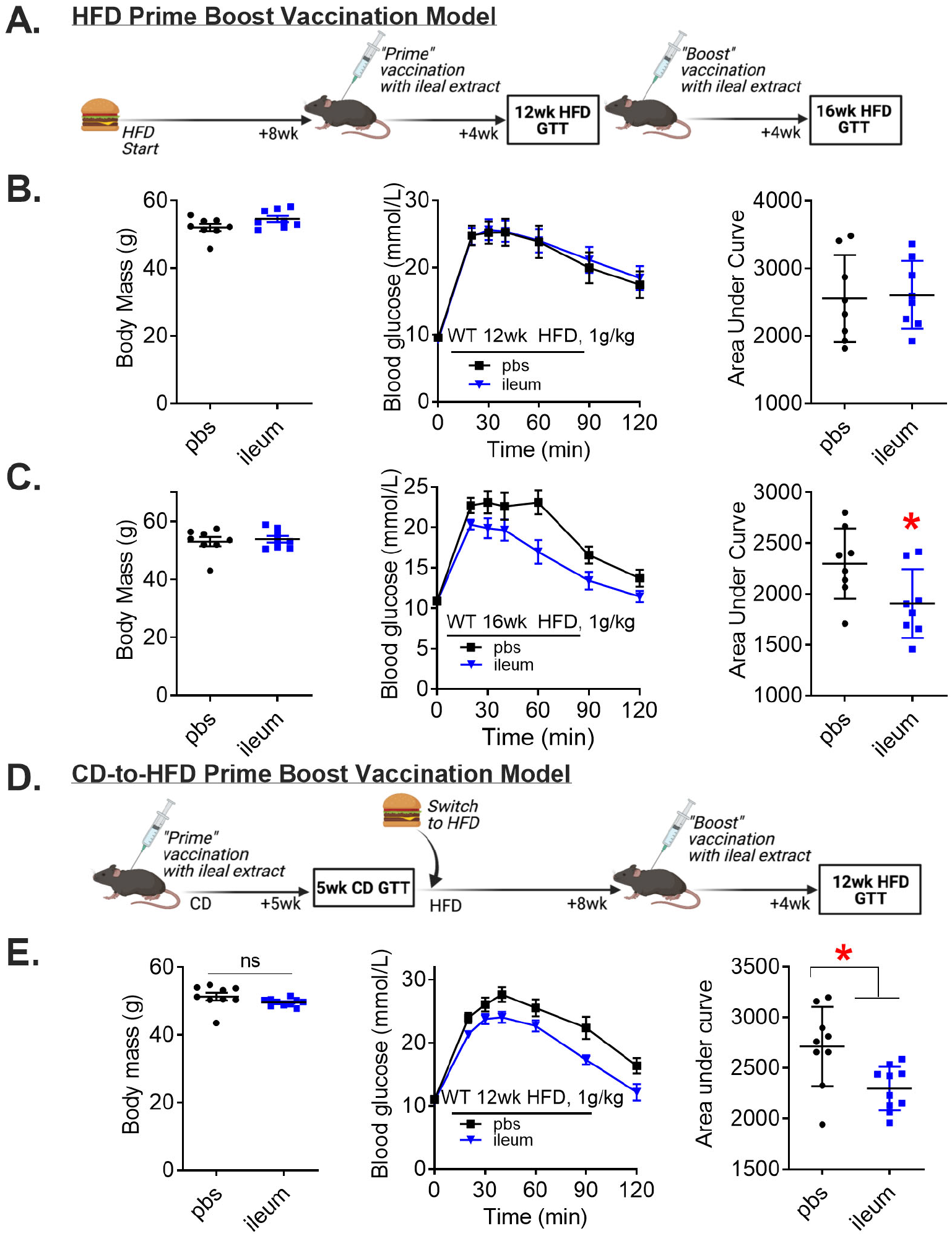
A prime-boost vaccination strategy is required to improve glucose control in obese male mice. **A)** Vaccination model during diet-induced obesity where mice were fed a HFD for 8 weeks before first ileal extract injection and glucose tolerance was assessed 4 weeks after the initial injection. A second ‘boost’ injection of ileal extract was delivered, and glucose tolerance was assessed 4 weeks after the second injection, which equated to a total of 16 weeks of HFD feeding. B) Body mass, blood glucose vs. time and quantified area under the curve during a glucose tolerance test (GTT) (1g/kg, i.p.) in 12wk HFD-fed WT male mice vaccinated once according to Fig 5C (n= 8). C) Body mass, blood glucose vs. time and quantified area under the curve during a GTT (1 g/kg, i.p.) in 16wk HFD-fed WT male mice vaccinated twice according to Fig 5C (n=8). D) Experimental vaccination model of given a “prime” injection of ileal extract during control diet (CD)-feeding in mice. Five weeks after the prime injection, mice were switched to a HFD for 8 weeks and then a second “boost” injection of ileal extract was delivered. Four weeks later, after 12 total weeks of HFD feeding, glucose tolerance was assessed. E) Body mass, blood glucose vs. time and quantified area under the curve during a GTT (1g/kg, i.p.) in 12wk HFD-fed WT male mice vaccinated twice according to Fig 5D (n=9-10). "Denotes significant different from control group injected with saline (pbs) determined by t-test (p < 0.05). Each symbol represents a mouse and other values shown are the mean +/− SEM.

### Muramyl dipeptide is not sufficient to improve blood glucose control when administered as a vaccination

We have previously shown that three consecutive injections of MDP requires NOD2 to lower blood glucose levels during a GTT in obese male and female mice (Cavallari et al., 2020a, 2017). Here, we found that *Nod2^−/−^* mice were refractory to changes in blood glucose after a vaccine-like ileal extract injection in mice (Figure 4C). Thus, we next sought to test if MDP was one component of the ileal extract that was sufficient to improve blood glucose control in our vaccination model (Figure 6A). Blood glucose levels were not altered five weeks after a single injection of MDP in CD-fed male mice (Figure 6B). Interestingly, a second injection of MDP after 8 weeks of HFD produced a significant increase in blood glucose levels during a GTT after mice were on a HFD for a total of 12 weeks (Figure 6C). These data indicate that MDP is not the pivotal ingredient responsible for improved blood glucose control caused by microbiota-based vaccination.

**Figure 6.**
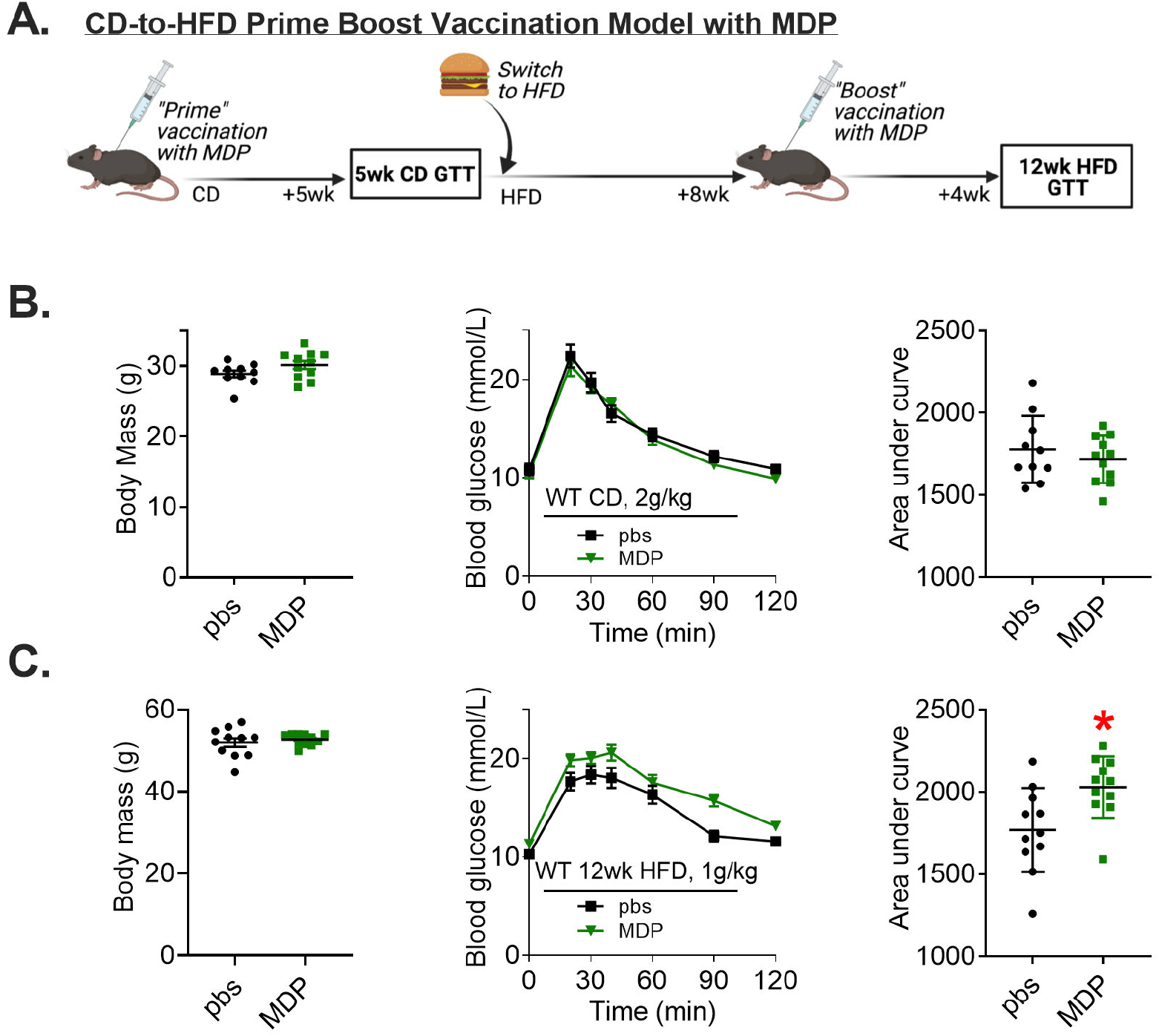
MDP is not sufficient to improve blood glucose control using a vaccination-style delivery. A) Vaccination model of CD-fed mice given a single injection of muramyl dipeptide (MDP). Glucose tolerance was assessed 5 weeks after the initial injection and mice were switched to a HFD for 8 weeks before a second injection of MDP was delivered. Four weeks after the second injection, after 12 total weeks of HFD feeding, glucose tolerance was assessed. B) Body mass, blood glucose vs. time and quantified area under the curve during a glucose tolerance teste (GTT) (2g/kg, i.p.) in control diet (CD)-fed WT male mice injected once with MDP according to Fig 7A (n=11). C) B) Body mass, blood glucose vs. time and quantified area under the curve during a GTT (1 g/kg, i.p.) in 12wk HFD-fed WT male mice injected twice with MDP according to Fig 7A (n=11). ‘Denotes significant different from control group injected with saline (pbs) determined by t-test (p < 0.05). Each symbol represents a mouse and other values shown are the mean +/− SEM.

### Microbiota-based vaccination increases bacterial extract-specific IgG in the ileum and alters the composition of the gut microbiome

Previous work showed that subcutaneous injection of a proximal gut bacterial extract caused a modest increase in total circulating immunoglobin-G (IgG) in mice (Pomié et al., 2016). However, total IgG levels in the blood do not necessarily capture compartmentalized adaptive immune or immunoglobin responses raised against specific antigens. An immune response after microbiota-based vaccination with intestinal extracts could produce a robust increase in IgG directed against the components in the bacterial extract, despite little or no difference in total IgG concentration. We analysed the serum and different gut segments of vaccinated mice using ELISA to detect antibodies specific to components of the intestinal extract 42 days after vaccination of CD-fed mice. Our results show that levels of ileal extract-specific IgG were unchanged in the serum of vaccinated mice (Figure 7A). However, we found an increase in the reciprocal endpoint dilution of IgG raised against the ileal extract antigens in the ileum, but not cecum or colon of vaccinated mice (Figure 7B-D; see Supplemental Figure 2 for full serial dilution curves). These data suggest that subcutaneous injection with a bacterial extract from the ileum elicits a specific B cell-mediated IgG response that is compartmentalized to the ileum of vaccinated mice.

**Figure 7.**
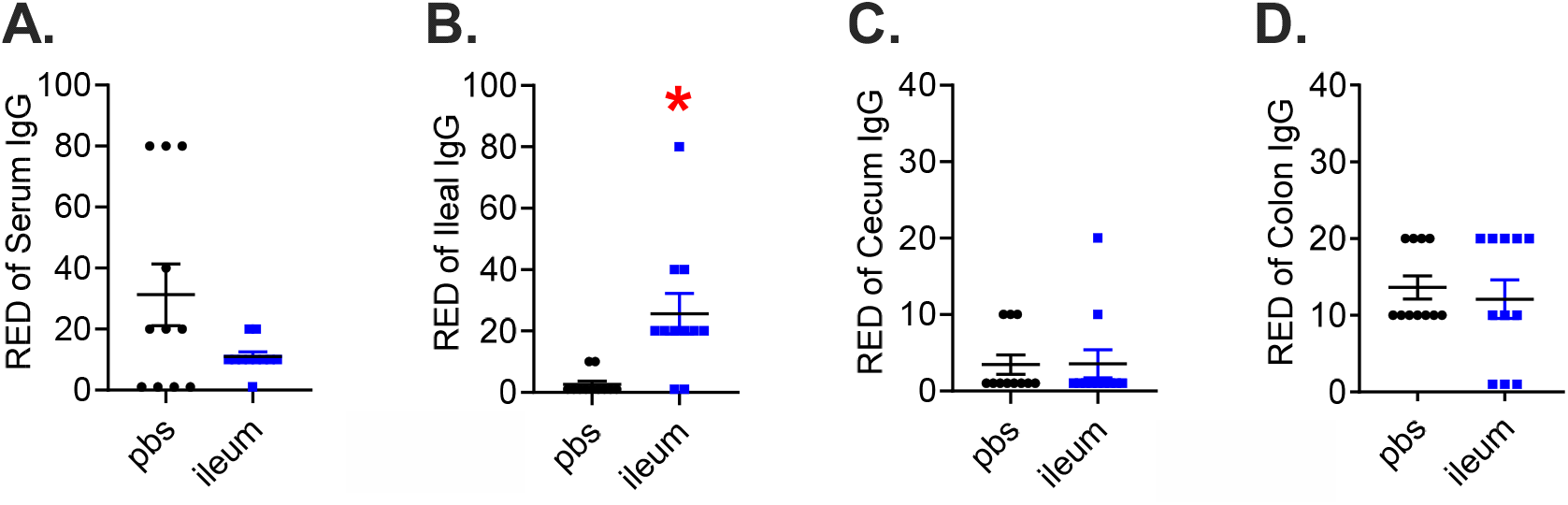
Quantification of extract-specific systemic and intestinal IgG levels in vaccinated mice. Reciprocal endpoint dilution (RED) of specific Immunoglobin-G (IgG) raised against ileal extract antigens in A) Serum, B) Ileum, C) Cecum, and D) Colon of ileal extract-vaccinated male mice (n=11/group for all tissues). *Denotes significant different from control group injected with saline (pbs) determined by t-test (p < 0.05). Each symbol represents a mouse and other values shown are the mean +/− SEM.

Based on these compartmentalized changes to intestinal immunity, we next sought to test if proximal or distal gut microbial communities were altered by vaccination. Amplicon-based sequencing was used to obtain the taxonomic bacterial profile in ileum and colon 42 days after vaccination in CD-fed male mice. PCoA scatterplots on Bray-Curtis distance revealed a significant degree of dissimilarity between the bacterial communities found in the ileum and colon of vaccinated mice (Figure 8A-B). This vaccination-driven clustering of bacterial populations was associated with changes in taxa within the phylum Firmicutes (Figure 8C-D). *Lachnospiraceae.NK4a136.group* was increased after ileal-vaccination, and this finding was consistent across the different gut niches analyzed (Figure 8E-F). After microbiota-based vaccination, the relative abundance of *Lachnospiraceae.NK4a136.group* increased to approximately 1/5^th^ of the entire pool of sequences in both the ileum and colon (Figure 8E-F). The relative abundance of *Clostridium.senso.strictu.1* and *Tenericutes* were decreased in the ileum and in the colon after the microbiota-based vaccination (Figure 8E-F). These data suggest that gut bacteria, and specifically *Lachnospiraceae.NK4a136.group,* are responsive to a microbiota-based vaccination and associated with increased ileal IgG responses and long-lasting improvements in blood glucose control.

**Figure 8.**
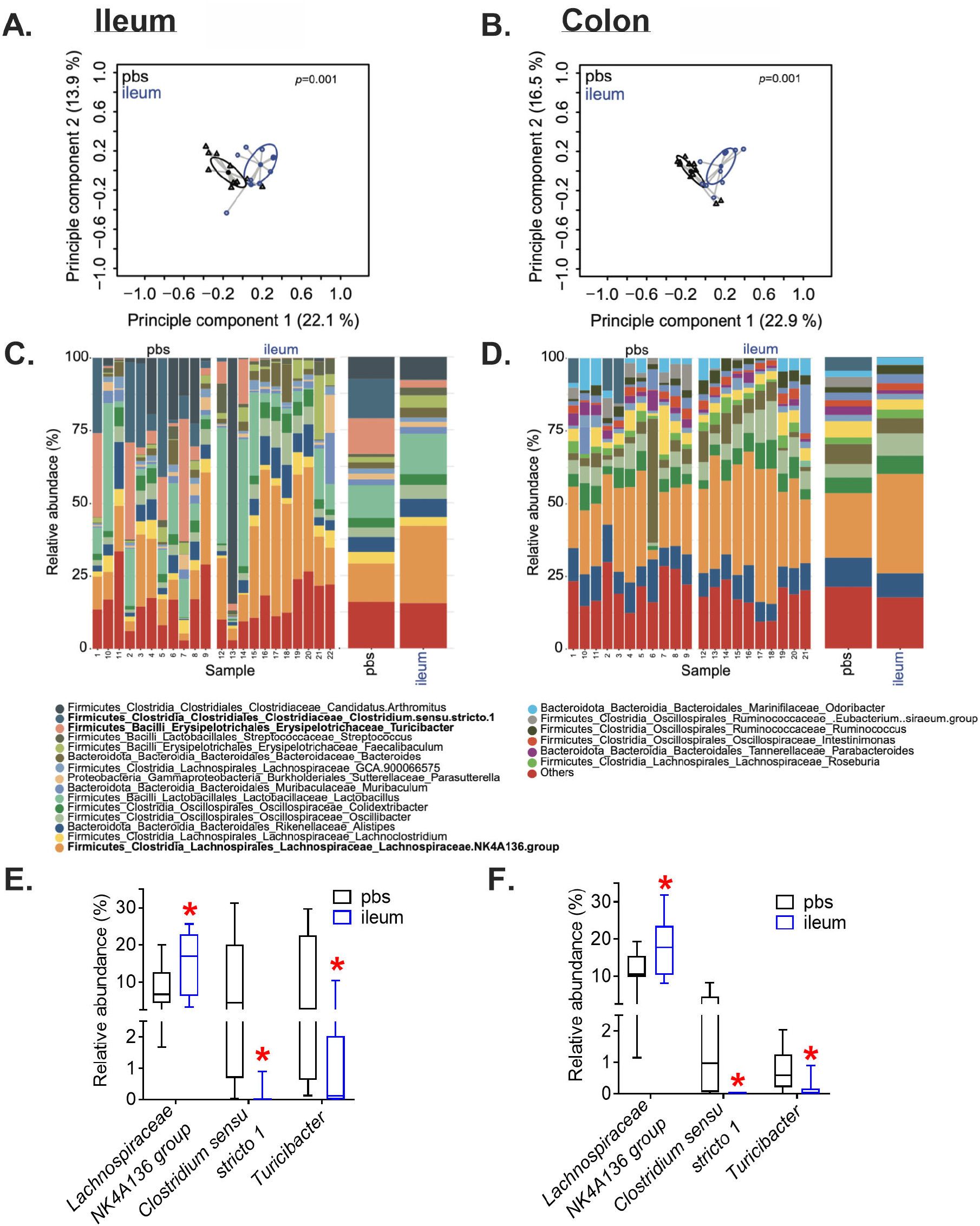
Microbiota-based vaccination alters ileal and colon gut microbiota composition. Principle coordinates analysis (PCoA) performed on Bray-Curtis distances in A) ileum and B) colon samples of CD-fed mice 42 days after vaccination with ileal extract (‘ileum’, blue circles) or pbs (black circles). Relative abundance of bacteria resolved to the phylum level in C) ileum and D) colon samples (with each bar representing an individual mouse) and the top three significant taxonomic differences in E) ileum and F) colon samples from CD-fed mice 42 days after vaccination with ileal extract (‘ileum’) or pbs (n= 10-11, p<0.05).

## Discussion

Commensal bacteria engage innate and adaptive immunity, which can alter host metabolism. The intestine has many barriers that limit how bacteria influence host immunity and metabolism. There is emerging evidence that bacteria can subvert the intestine and that T2D dictates the type and amount of bacteria that can penetrate into metabolic tissues that control blood glucose (Anhê et al., 2020). Smaller molecules such as bacterial cell wall components and metabolites can also translocate across the intestinal barrier and engage immune response in host tissues that influence blood glucose control (Cani et al., 2008; Chan et al., 2017). For example, microbiota-derived bacterial cell wall components such as meso-DAP-containing muropeptides and LPS promote inflammation, lipolysis and insulin resistance (Cani et al., 2008; Chan et al., 2017; Chi et al., 2014; Schertzer et al., 2011). Conversely, bacterial MDP promotes immune tolerance and insulin sensitivity (Cavallari et al., 2020a, 2017). How these opposing bacterial factors, particularly in a complex microbial extract that contains a plethora of bacterial metabolites and components, alter blood glucose control is unknown. One previous report showed that a single injection of a proximal gut bacterial extract improved blood glucose control in mice, and this was linked to a systemic adaptive immune response (Pomié et al., 2016). Our results show that a microbiota-based vaccination generates a compartmentalized adaptive immune response where IgG directed against the antigens within the bacterial extract is increased in the proximal gut of mice receiving the injection. Further, our results show that microbiota-based vaccination improvements in blood glucose control requires a NOD2-RIPK2-mediated innate immune response in the mice injected with bacterial extracts. Therefore, our results define the innate immune bacterial detection pathway and compartmentalized adaptive immune responses that are associated with improvements in blood glucose after injection of postbiotic bacterial components in a proximal gut bacterial extract.

Despite this need for NOD2 to propagate the glucose effects of a microbiota-based vaccination, a single subcutaneous injection of the well-known NOD2 ligand, MDP, was not sufficient to generate a long-lasting improvement in blood glucose control. It is possible that unclassified NOD2 ligands (beyond MDP) could be factors in bacterial extracts that improve blood glucose control. It is plausible that unknown bacterial factors within the bacterial extract engage innate immune receptors beyond NOD2, where the interaction of multiple innate immune pathways require NOD2 immunity to drive an IgG-mediated adaptive immune response in the proximal gut, and subsequently changes in the composition intestinal microbiome and glycemia. The identities of these other bacterial factors are not clear, but other bacterial components that engage separate innate immune responses could participate in an integrated response that alters blood glucose. For example, injection of flagellin can improve blood glucose control in mice by engaging TLR5 and adaptive immune responses that can protect against aspects of inflammatory and metabolic diseases (Tran et al., 2019). It is beyond the scope of the current paper to test all the possible bacterial factors that could interact to elicit long-lasting changes in blood glucose control. However, it is noteworthy that injection of a specific concentration of proximal gut bacterial extracts was required for improvements on blood glucose control, whereas extracts derived from the cecum or large intestine (or feces) did not alter blood glucose. This could be related to the type of bacteria, type of bacterial components, or concentration of bacteria, which are much higher in the distal gut. Determining the concentration and gut-segment-dependent effects may help elucidate the identity of postbiotic components that contribute to improved blood glucose control.

We have resolved that NOD2 is a critical component of innate immunity that mediates improved glycemic responses to microbiota-based immunization in male mice. We also found that deletion of NOD1 or RIPK2 worsened blood glucose control after microbiota-based vaccination. RIPK2 is the common adapter for both NOD1 and NOD2 and our data show that the NOD1-RIPK2 pathway protects against dysglycemia after subcutaneous injection of bacterial extracts. How or why this occurs is not known, it is possible that augmented TLR4 immune and lipid metabolism responses that can occur in the absence of RIPK2 could be mediators of dysglycemia (Levin et al., 2011). We have consistently shown that repeated MDP engagement of NOD2 is an innate immune component that improves blood glucose control (Cavallari et al., 2020b, 2020a, 2017). Therefore, based on our current data, we propose a new model where the proximal gut luminal contents contain a complex milieu of postbiotic components derived from the commensal microbiota that can either raise or lower glycemia and NOD1-RIPK2 versus NOD2-RIPK2 signalling dictates the immune response and subsequent net effects on glycemic control.

We found that changes in blood glucose and the compartmentalized intestinal IgG response were associated with changes in resident ileal and colon microbial communities in vaccinated mice. Specifically, the abundance of *Lachnospiraceae.NK4a136.group* was significantly increased in the ileum and colon of vaccinated mice.

*Lachnospiraceae.NK4a136.group* is a potential butyrate-producer associated with metabolic fitness and gut homeostasis (Hu et al., 2019; Ma et al., 2020) and probiotic activity in mice (Wu et al., 2020). We found that *Lachnospiraceae.NK4a136.group* constituted ~20% of the sequenced bacterial taxa in vaccinated mice, which makes it a strong candidate for future testing related to glycemic control. Although *Lachnospiraceae.NK4a136.group* was increased in both the ileum and colon of vaccinated mice, bacterial vaccination only increased IgG responses in the ileum. Hence, our data do not directly link compartmentalized adaptive immunity to changes in the composition of the microbiota within each gut segment. It is plausible that microbial vaccination-induced increases in proximal gut IgG influence the local gut microbiota in the ileum, where compositional changes in the microbiome carryover into the distal gut (i.e., colon), but directionality in this host-microbe relationship has not yet been established. Although most of our experiments were focussed on male mice, we did find similar improvements in glycemic control in vaccinated female mice, suggesting a sex-independent effect on glycemia. However, a more thorough analysis of any sex-specific responses to microbiota-based vaccination across the life-course is warranted, given the sex- and age-related differences in gut microbiota composition, immunity and metabolism (Org et al., 2016).

In summary, we found that a simple microbiota-based vaccination procedure using subcutaneous injection of a proximal gut bacterial extract, elicits a proximal gut IgG response directed against the antigens in the bacterial extract, and promotes improvements in blood glucose control in male and female mice. The proximal gut extract required bacteria and engaged NOD2-RIPK2 immunity to improve blood glucose control in male mice. Obese male mice had to receive prime and boost injections for improvements in blood glucose control. There is a lot of interest in manipulating the gut microbiota to improve metabolism during obesity. Given that some probiotic and FMT strategies aimed at combating metabolic disease have limitations that rely on live bacteria establishing a transient niche in the gut, investigation of postbiotics that promote long-term changes in blood glucose is warranted. Our results show that the net effect of injecting a specific concentration of postbiotics derived from the proximal gut produces a long-lasting improvement in blood glucose control in mice.

## Abbreviations

T2D: Type 2 Diabetes
FMT: fecal microbial transfer
LPS: lipopolysaccharide
meso-DAP: meso-diaminopimelic acid
MDP: muramyl dipeptide
CD: control diet
HFD: high fat diet
GTT: glucose tolerance test
ITT: insulin tolerance test
OGSIS: oral glucose-stimulated insulin secretion
PGN: peptidoglycan
PRR: pattern recognition receptor
IgG: immunoglobin-G
RED: reciprocal endpoint dilution

---

**Supplemental Figure 1.**
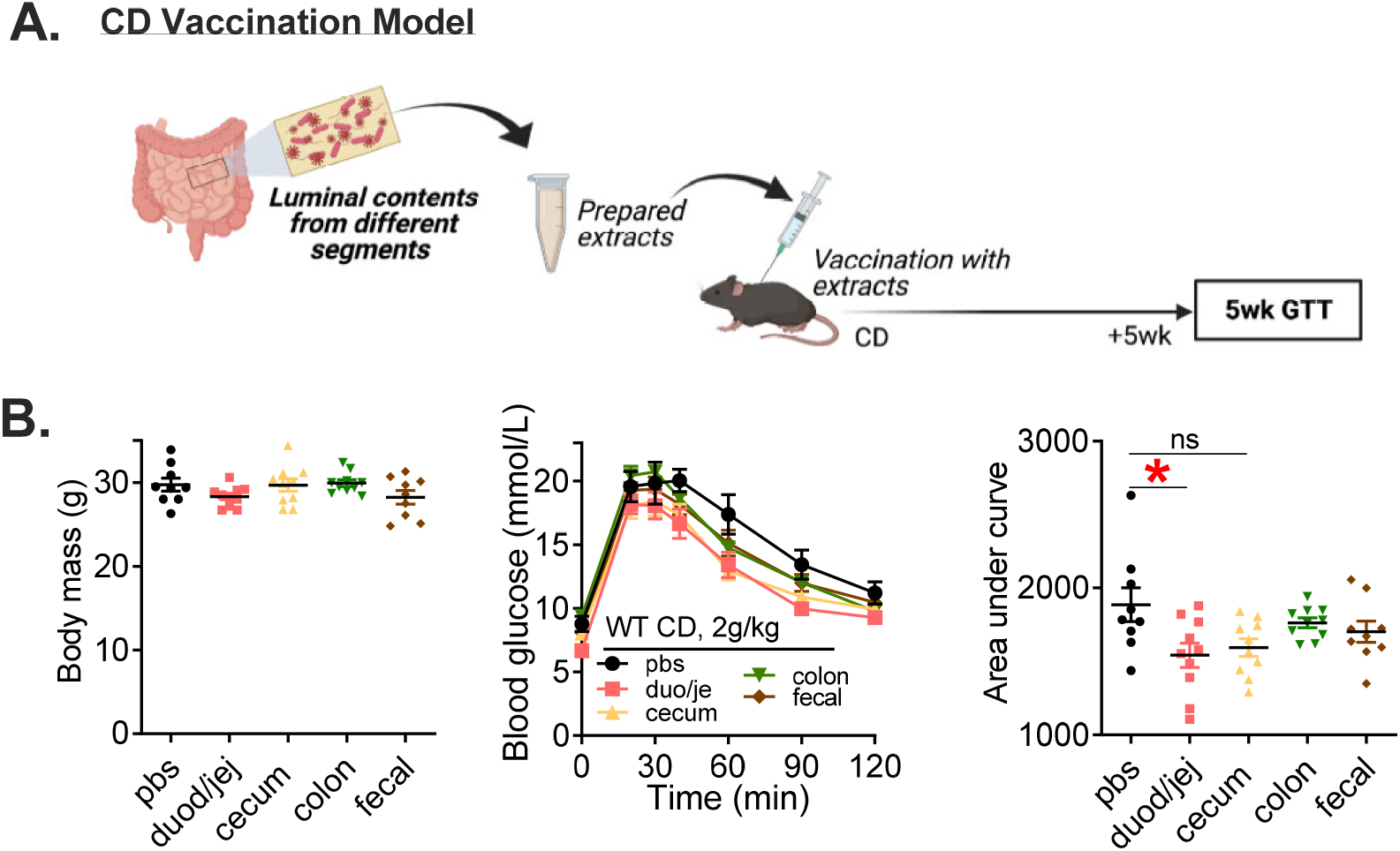
Microbiota-based vaccination with proximal gut extracts improves blood glucose control in male mice. A) Vaccination model of control diet (CD)-fed male mice given a single subcutaneous injection with extracts prepared from different intestinal segments (dudodenum/jejenum, cecum, colon) or from a homogenized fecal pellet (‘fecal’) from male mice. Glucose tolerance was assessed 5 weeks after the vaccination event. B) Body mass. Blood glucose vs. time and quantified area under the curve during a glucose tolerance test (GTT) (2g/kg, i.p.) in CD-fed WT male mice 5 weeks after vaccination (5000x dilution was used for all extracts; n=9-10). *Denotes significant different from control group injected with saline (pbs) determined by one-way ANOVA (p < 0.05). Each symbol represents a mouse and other values shown are the mean +/− SEM.

**Supplemental Figure 2.**
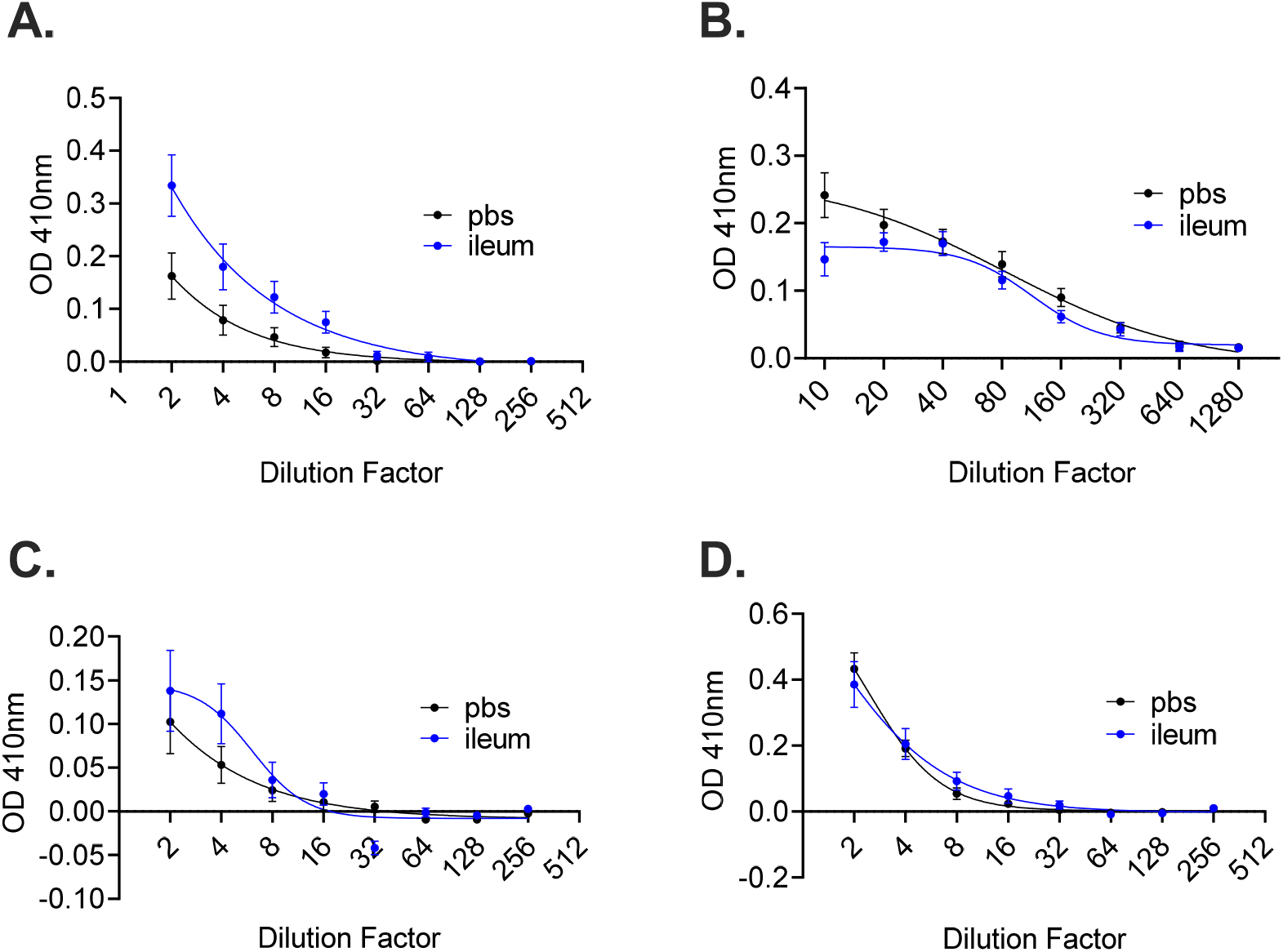
Quantification of IgG levels directed against the extract antigens in vaccinated mice. Serial dilutions of A) serum, B) ileum, C) cecum, and D) colon samples for detection of specific Immunoglobin-G (IgG) raised against ileal extract antigens in ileal extract-vaccinated mice (n=11/group for all tissues). IgG was detected in mice 42 days after a single subcutaneous injection of ileal extract (5000 x dilution) or saline (pbs). Values shown are the mean +/− SEM.

